# Measuring brain beats: cardiac-aligned fast fMRI signals

**DOI:** 10.1101/2022.02.18.480957

**Authors:** Dora Hermes, Hua Wu, Adam B. Kerr, Brian Wandell

## Abstract

Blood and cerebrospinal fluid (CSF) pulse and flow throughout the brain, driven by the cardiac cycle. These fluid dynamics, which are essential to healthy brain function, are characterized by several noninvasive magnetic resonance imaging (MRI) methods. Recent developments in fast MRI, specifically simultaneous multislice (SMS) acquisition methods, provide a new opportunity to rapidly and broadly assess cardiac-driven flow, including CSF spaces, surface vessels and parenchymal vessels. We use these techniques to assess blood and CSF flow dynamics in brief (3.5 minute) scans on a conventional 3T MRI scanner. Cardiac pulses are measured with a photoplethysmograph (PPG) on the index finger, along with fMRI signals in the brain. We retrospectively analyze the fMRI signals gated to the heart beat. Highly reliable cardiac-gated fMRI temporal signals are observed in CSF and blood on the timescale of one heartbeat (test-retest reliability within subjects R^2^>0.50). Cardiac pulsations with a local minimum following systole are observed in blood vessels, with earlier extrema in the carotid and basilar arteries and in branches of the anterior, posterior and middle cerebral arteries and extrema ∼200 ms later in the superior sagittal, transverse and straight sinuses. CSF spaces in the ventricles and subarachnoid space showed cardiac pulsations with a local maximum following systole instead. Similar responses are observed, with less temporal detail, in slower resting state scans with slice timing retrospectively aligned to the cardiac pulse in the same manner. The SMS measurements rapidly, noninvasively and reliably sample brain-wide fMRI signal pulsations aligned to the heartbeat. The measurements estimate the amplitude and phase of cardiac driven fMRI pulsations in the CSF relative to those in the arteries, which is thought to be an estimate of the local intracranial impedance. Cardiac aligned fMRI signals can provide new insights about fluid dynamics or diagnostics for diseases where these dynamics are important.

## 1. Introduction

It is important to understand the connection between the brain’s cardiovascular health and cognition. Preclinical studies in aging rodents show that arterial changes may precede cognitive decline (Nation et al., 2019). In COVID-19, which is primarily thought to affect cardiovascular function, long cognitive symptoms have been observed (Boldrini, Canoll, & Klein, 2021). The ability to assess cardiovascular efficacy within spatially resolved brain regions, and to connect this assessment to behavior, is also an important direction in MRI research (Wåhlin & Nyberg, 2019). A comprehensive characterization of the status of the brain’s fluid dynamics in individual participants can become an important diagnostic tool for cognitive and affective neuroscience. Including this assessment as part of a typical cognitive neuroscience functional MRI (fMRI) experiment may also help to clarify a source of differences between experimental participants.

Blood and cerebrospinal fluid (CSF) pulse through the brain, in synchrony with the cardiac cycle (Womersley, 1955). About each second a heartbeat produces a pressure wave; blood from the heart traverses the arterial to venous network during multiple heartbeats (Mihara et al., 2003). Various noninvasive MR methods provide specific information about the structure and function of the neurovascular system. MR angiography sequences provide insight into the integrity of the vascular anatomy either by the use of intravenous contrast to enhance blood signal, by designing sequences to manipulate the blood signal relative to be much darker than other tissue (black-blood), much brighter (bright-blood) (Nakao et al., 2018), or by introducing flow-sensitive encoding (phase-contrast) (Pelc, Bernstein, Shimakawa, & Glover, 1991). Arterial spin labeling quantitatively and noninvasively estimates brain perfusion with arterial blood water (Bambach, Smith, Morris, Campeau, & Ho, 2020; Telischak, Detre, & Zaharchuk, 2015).

Other measurements can be cardiac-gated to assess the dynamic properties of the cardiac pulse. Acquiring one slice at various times after scanning software detects a peak in the heartbeat in real time. In MR elastography a vibrating source is used during scanning to deform the tissue and estimate its stiffness (Glaser, Manduca, & Ehman, 2012; Kruse et al., 2008; Manduca et al., 2001). A cardiac-gated MR elastography sequence showed that cerebral vascular compliance, the degree to which tissue absorbs the cardiac pulse, decreases in older individuals (Schrank et al., 2020). Cardiac gated cine MRI methods have also been used to measure dynamic changes in the MR signal (Curtis & Cheng, 2021) and assess the velocity of blood flow in a few slices perpendicular to the major cerebral arteries or the cerebral aqueduct (Enzmann, Ross, Marks, & Pelc, 1994).

These methods are not part of the standard suite of tools used in cognitive neuroscience despite the fact that the spatial distribution of the pulsatile dynamics in blood and CSF spaces change in multiple neurological and neuropsychiatric diseases (Nation et al., 2019; Wåhlin & Nyberg, 2019). For example, aging and Alzheimer’s disease (AD) alter the rigidity of the arteries, impacting the shape of the flow pulsations. Transcranial doppler ultrasonography has revealed that increased vascular rigidity increases the pulsatility of the blood flow in the circle of Willis (Roher et al., 2006). Increases in pulsatility with aging may also occur deeper in the microvasculature, which may result in lesions in many brain areas (for reviews see (Tsvetanov, Henson, & Rowe, 2021; Wåhlin & Nyberg, 2019). In addition, aging can result in increases in the variability of the fMRI signal (Tuovinen et al., 2020). Variability in resting state fMRI signals have been related to cardiovascular dynamics (Bayrak et al., 2021; Chang, Cunningham, & Glover, 2009; Chen et al., 2020; Shmueli et al., 2007). Cardiac pulsations can thus affect vascular changes in tissue where studies of perception, action and cognition measure fMRI responses.

The method we describe here builds on these technologies but focuses on the amplitude and phase of the dynamic fMRI signal. Recent development of simultaneous multislice (SMS) methods permits investigators to sample the fMRI signal of the whole brain multiple times within a single cardiac cycle (Larkman et al., 2001; Setsompop et al., 2012). Individual slices can be acquired with a duration of less than 50 ms, and multiple groups of slices are obtained within a single cardiac cycle. Studies measuring this fMRI signal have shown that changes in heart rate can result in changes in the fMRI signal that span several heart beats (Chang et al., 2009). In single slice acquisitions measured at 7T, cardiac pulsations could be measured in blood vessels and CSF spaces (Bianciardi et al., 2016; Viessmann, Möller, & Jezzard, 2017). We measure whole-brain fMRI signals and assess how well these signals can be tracked at 3T within a single heartbeat. By aggregating measurements over multiple cardiac cycles and retrospectively aligning the measurements to the cardiac cycle onset, we obtain a noninvasive assessment of the pressure wave and flow of the vasculature and the CSF in many locations across the cranium.

## 2. Methods

In this work we retrospectively align fMRI signals measured at 3T to the cardiac pulse using a widely used SMS protocol. In this sequence, forty slices are measured in five groups, each group comprising eight separated planes. The data in a single group of planes is acquired simultaneously over a 50 ms interval; the entire brain - all five groups - is measured every 250 ms. The typical cardiac cycle duration is about 1000 ms. The measurements cover the whole-brain and provide spatially-resolved information of 4 mm isotropic voxels. We compared these responses to fMRI responses measured during a typical resting state fMRI scan (TR = 2s, 240 volumes). Even in a single-subject, the cardiac-aligned time series modulations can be very reliable. The most reliable cardiac aligned modulations are located near the principal arteries and veins, portions of the CSF, and certain gray matter regions. Blood vessels showed local minima around the time of the PPG peak while CSF spaces showed local maxima. These response features were consistent across the SMS and slow fMRI sequence. This paper describes the measurements, analytical methods and signals in healthy subjects.

### 2.1 Subjects and IRB statement

Data from ten healthy subjects are analyzed in the study. The study was approved by the Stanford University Institutional Review Board and all subjects provided informed consent to participate in the study. Five subjects were recruited specifically for measurements using fast SMS methods. One of these subjects was scanned twice, separated by a three-year interval. We analyzed additional data from five subjects who were part of a different study that obtained resting state data as a control condition. The acquisition parameters for both sets of subjects are described below.

### 2.2 Anatomical magnetic resonance imaging

All subjects were scanned on a 3T General Electric MRI 750 scanner at the Stanford Center for Cognitive and Neurobiological Imaging. In order to localize different anatomical gray matter regions and CSF spaces we acquired a T1-weighted anatomical image (1×1×1 mm voxels). We segmented the T1 weighted scan using Freesurfer (http://surfer.nmr.mgh.harvard.edu/, (Fischl, 2012)). In order to understand the effects of the arterial pulsations, we assigned each gray matter region from the Desikan-Killiany Atlas (Desikan et al., 2006) to a ‘distance’ level varying in 4 steps from closest to furthest from one of the main three arterial branches: the anterior cerebral artery (ACA), middle cerebral artery (MCA) and posterior cerebral artery (PCA). To map veins, we collected an MR venogram (0.43 × 0.43 × 2 mm voxels) and thresholding this image allowed us to select the superior sagittal sinus.

In order to compare the location of reliable cardiac averaged responses across subjects, the non-linear transformation to MNI152 space was calculated based on the T1 scan using Unified Segmentation in SPM12 (Ashburner & Friston, 2005).

### 2.3 Cardiac cycle measurements using PPG

To estimate the cardiac cycle a pulse oximeter was attached to a finger and the photoplethysmograph (PPG) signal was measured during MRI scanning (Figure 1A). The peak in the autocorrelation of the PPG signal revealed the heart rate, such that peak detection could be done with a minimum peak distance of 70% of the heartbeat cycle. Before detecting the peaks, PPG data were low pass filtered at 5Hz using a 3^rd^ order Butterworth filter in two directions. This allowed the detection of the peaks in the PPG signal (known to be related to systole) that occurred around every heartbeat cycle. We found this method to be overall more reliable compared to the peaks detected by the scanner software (Supplementary Figure 1).

**Figure 1:**
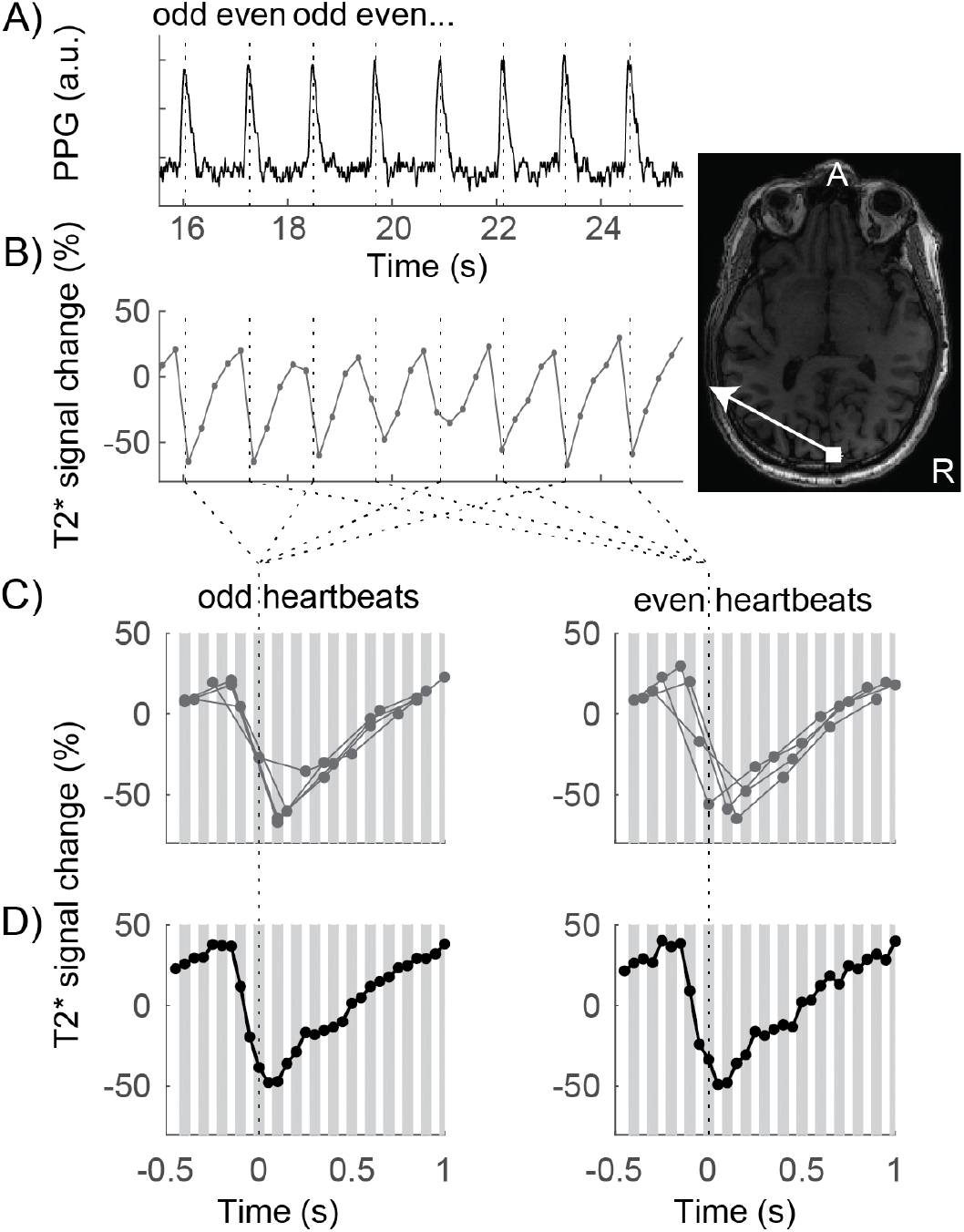
Temporal alignment of MRI measurements to heartbeat. **A)** The PPG signal was measured with a pulse oximeter. Peaks in the signal were detected (dotted lines). **B)** The fMRI signal from a voxel near the superior sagittal sinus was measured every 250 ms, for a duration of 50 ms. **C)** Signals were aligned to even and odd heartbeat peaks, gray lines indicate the 50 ms within which each time-point was measured. This example shows the signals from the eight heartbeats in B) **D)** The signals averaged for all even and odd heartbeats form a smooth curve of the cardiac-gated response, sampled every 50 ms.

### 2.4 Functional magnetic resonance imaging

In five subjects, we acquired simultaneous multislice, fMRI measurements to map brain wide cardiac-gated fMRI signal variations. Whole-brain fMRI data were acquired using several EPI sequences with whole brain coverage and simultaneous multislice (SMS, sometimes also called multiband or hyperband) (Larkman et al., 2001; Setsompop et al., 2012). First, an SMS sequence was used with 4mm isotropic voxels, a flip angle of 48 degrees, TR = 250ms, TE = 11.6ms, FOV = 224×224 and 40 slices with multiband factor 8. This resulted in a slice acquisition time of 50 ms. A total of 878 volumes were acquired within a total scan duration of 220 sec.

In five additional subjects, we analyzed data from a typical resting state fMRI study (Hack et al., 2021) where PPG signals were acquired. A whole brain EPI sequence was used with 3mm isotropic voxels, a flip angle of 77 degrees, TR = 2s, TE = 27.5ms, FOV = 216×216 and 45 slices. A total of 240 volumes were acquired within a total scan duration of 480 sec.

Every slice is acquired at a different time with respect to the cardiac pulse, and we wanted to preserve the 50ms (SMS) and 44.4ms (resting state) sampling per slice. It was therefore essential to preserve the measured voxel time series and not perform slice timing correction. The fMRI data were inspected for motion and no data were excluded for motion. The linear transformation matrix to spatially align the fMRI with the anatomical data was calculated to align the functional volumes.

### 2.5 Calculation of cardiac aligned responses

We calculated cardiac averaged responses as follows. The first three volumes were removed. The remaining time measurements were transformed to units of percent modulation and linear trends were removed (Figure 1B). For every voxel in each slice, the slice time was noted with respect to the heartbeat peaks on the PPG signal measured on a finger (Figure 1C). Each slice was measured within a 50 ms temporal window, and the average PPG-peak-aligned signal was calculated pooling from the entire acquisition (Figure 1D). Given that heartbeats are spaced about 1 s duration, there are about 20 sample windows between heartbeats.

To calculate reliability of these cardiac-gated responses, we averaged signals separately from the even and odd PPG peaks (Figure 1D). We then calculated the coefficient of determination between even and odd heartbeats. The coefficient of determination (R^2^) is a measure of the reliability of the cardiac-gated time series.

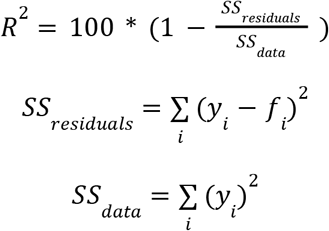

where *y*_*i*_ is the even response amplitude and *f*_*i*_ is the odd response amplitude for time point *i*. Note that *R*^*2*^ is defined here with respect to zero, rather than with respect to the mean response, to avoid the arbitrariness of the mean.

### 2.6 Modeling of cardiac averaged responses

Capturing the amplitude and temporal delay of the cardiac pulsatility in different cranial compartments may be important to further understand disease mechanisms. In order to quantify the temporal delay and amplitude of the responses, we developed a parameterized model of these waveforms using singular value decomposition based on the SMS time series time series data. The amplitude of the temporal waveforms of the cardiac averaged responses in each voxel was set to the *R*^*2*^, such that the largest responses entering the SVD are the ones that are the most reliable across the cardiac cycle, and the smallest responses were those that are the least reliable. This results in a matrix *M* of dimensions voxels by time, which was decomposed using singular value decomposition: *M = U*Σ*V*\*. The columns of *U* contain the eigenvectors as a function of time, and *V* contains the spatial weighting for these vectors across the brain. The SVD was performed on half the data from the odd heart beats. We then calculated the number of components that explained over 70% of variance in the other half of the data from the even heart beats.

We used the following strategy to develop a model across the subjects. The principal components were calculated in each individual subject as a function of time and these were then resampled across the heartbeat cycle to create standardized responses as a function of heartbeat cycle. A second SVD was then done on these components, leaving one subject out for cross-validation, resulting in a set of canonical principal components. A weighted combination of these canonical components could then be used to predict the cardiac aligned responses from the odd heart beats in the left out subject (*Y =* β_1_ *pc*_1_ + … + β_*n*_ *pc*_*n*_ + *ϵ*). We then calculated the relative root mean squared error (*R*_*rmse*_), for each voxel. The *R*_*rmse*_ describes how well the model (fitted on the training subjects) explained the cardiac averaged even heartbeat responses in the test subject compared to the test-retest reliability and is defined as 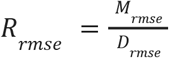, where *M*_*rmse*_ is the model prediction error and *D*_*rmse*_ is the data prediction error between even and odd responses. If *R*_*rmse*_ < 1, the model fitted on the other subjects predicts the data better compared to the within subject test-retest reliability. If a model perfectly predicts a signal with zero-mean Gaussian noise and standard deviation σ, the expected value of the  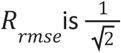.

### 2.7 Data and code availability

All our methods are open-source and shared on GitHub (LINK). Data are available in BIDS format on OpenNeuro.org under (LINK).

## 3. Results

In order to map brain-wide cardiac pulsations, we used a rapid fMRI acquisition and retrospectively calculated cardiac aligned responses. Figure 2A shows brain regions that have reliable responses at the frequency of the cardiac cycle; not all voxels have reliable cardiac aligned responses. First, voxels close to the sagittal and straight sinus show reliable responses with a local minimum around the same time as the peak of the PPG curve (time zero for cardiac aligned responses) (Figure 2B-C). Second, voxels close to the posterior cerebral artery, basilar artery and anterior cerebral artery similarly show reliable responses with a local minimum 0.1-0.2 sec earlier compared to the PPG peak (Figure 2F-H). Third, voxels near CSF regions such as subarachnoid spaces and the lateral ventricle show reliable responses with a local maximum around the same time as the PPG peak (Figure 2D-E). Importantly, not all brain regions show large cardiac pulsations, such as an area in the posterior gray matter that is distant from arteries, veins and CSF spaces (Figure 2I).

**Figure 2.**
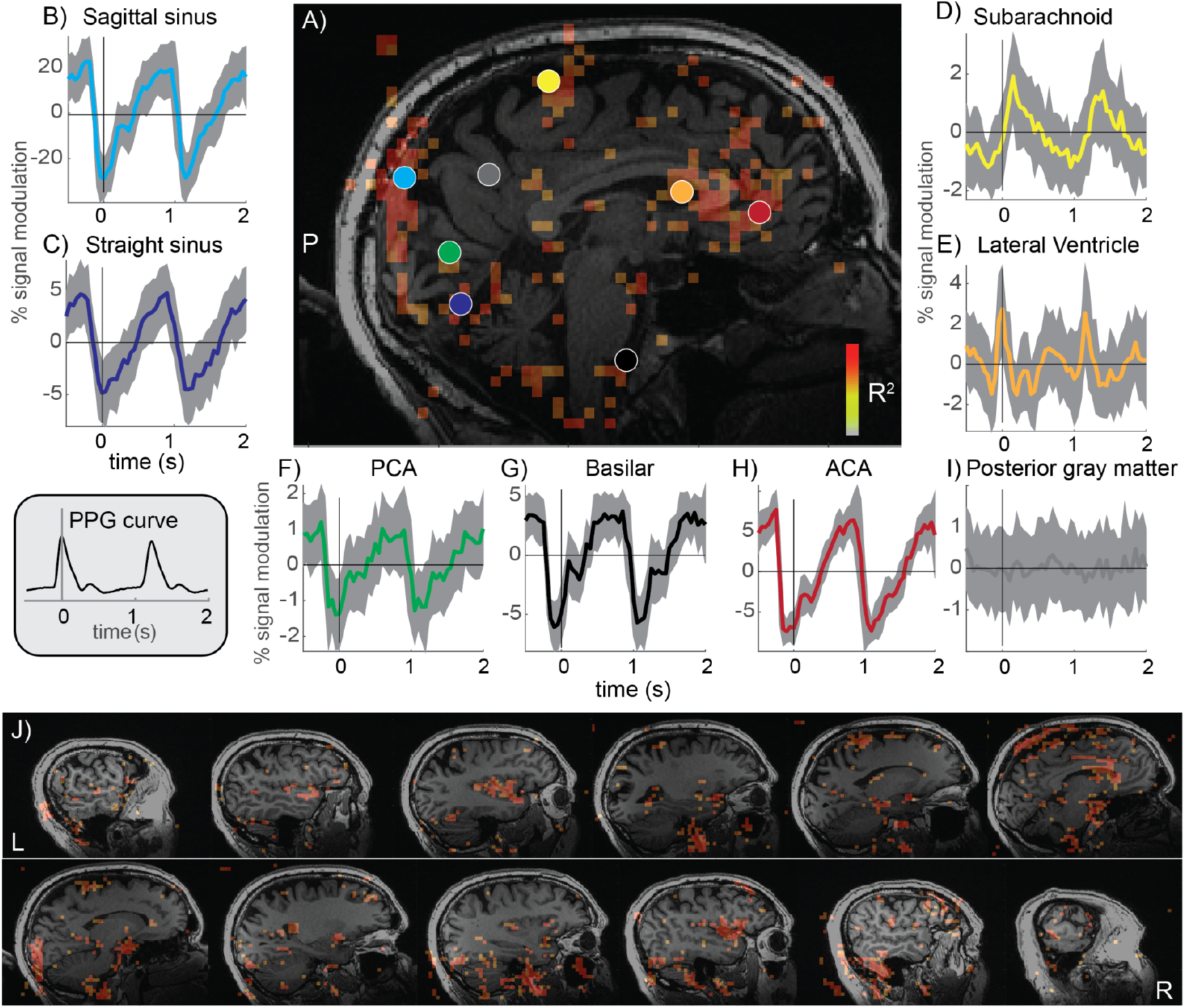
Reliable cardiac-aligned responses in a single subject. **A)** Sagittal slice showing voxels that had highly reliable cardiac aligned responses (R^2^>0.8). The colored circles indicate locations of the responses in panels B-I. The average (+/- standard deviation) cardiac aligned time series are plotted; zero indicates the peak in the PPG responses (gray inset). Cardiac aligned responses are shown for the sagittal sinus **(B)**, straight sinus **(C)**, subarachnoid **(D)**, lateral ventricle **(E)**, posterior cerebral artery **(F)**, basilar artery **(G)**, anterior cerebral artery **(H)** and posterior gray matter **(I). J)** Set of sagittal slices in the same subject shows the spatial distribution of the reliable cardiac aligned responses.

### 3.1 Regions with reliable cardiac aligned responses

We then consider which areas show reliable cardiac pulsations across the group of subjects, as large deviations from the group may be related to pathology (Rajna et al., 2021). Across the group of subjects, we find that some areas consistently show reliable cardiac pulsations whereas other areas do not. To look at the locations that are most consistent across subjects, the maps of reliable cardiac pulsations (Figure 2J) are converted to MNI152 space and averaged (Figure 3). In the average across the subjects it can be seen that reliable cardiac pulsations are observed near the ventricles, in areas close to the main three cerebral arteries (ACA, MCA, PCA), and in areas near large veins such as the superior sagittal sinus. Other regions (e.g., white matter) are consistent in showing no cardiac aligned modulation.

**Figure 3.**
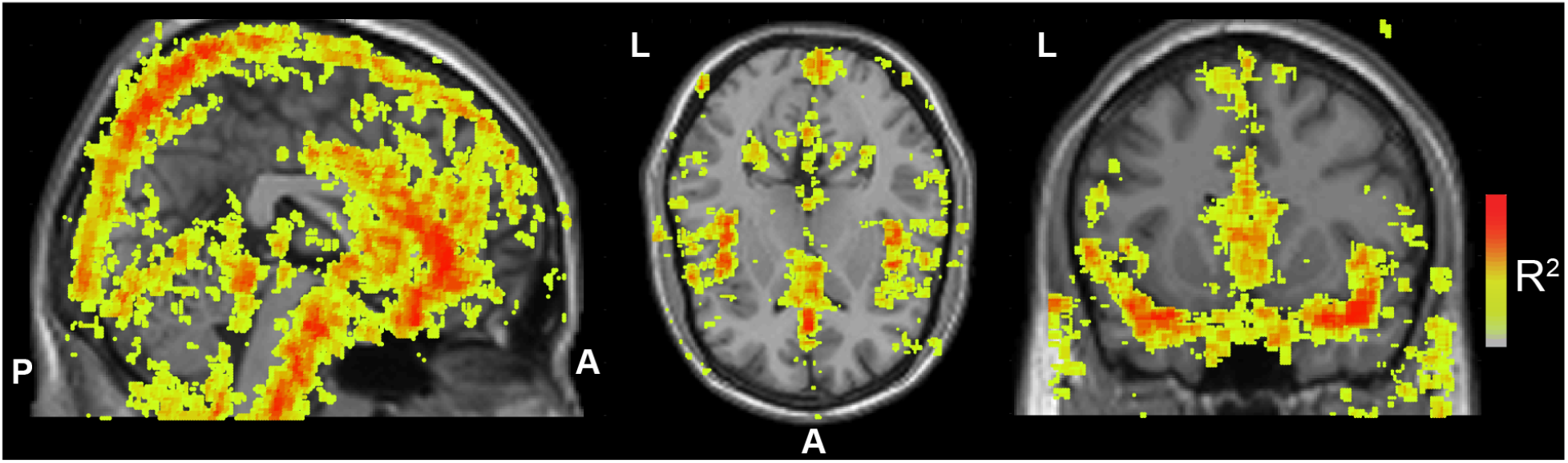
Reliable cardiac aligned response modulations across all subjects. For all subjects, maps of the reliability of the cardiac aligned responses represented in MNI152 space (without smoothing). Colors overlaid on a standard MNI brain indicate the average R^2^ across subjects for a sagittal, axial and coronal slice. Maps are thresholded and only voxels with an average R^2^>0.3 are shown to highlight those areas with reliable modulations.

### 3.2 Regional characterization of the cardiac aligned response shape

To understand whether the waveform of the cardiac aligned responses seen in Figure 1 is reliable across subjects, we segmented the T1 anatomical scan into several anatomical regions (Figure 4A). Figure 4B shows that reliable cardiac signals are observed in the raw fMRI signal from voxels located at the anterior cingulate (a brain region close to the anterior cerebral artery), the lateral ventricles and the superior sagittal sinus. The anterior cingulate and sagittal sinus waveforms have a local minimum at the time of the PPG peak. The lateral ventricles, however, show local maxima at the same time. These response shapes are highly robust across the subjects (Figure 4D).

**Figure 4.**
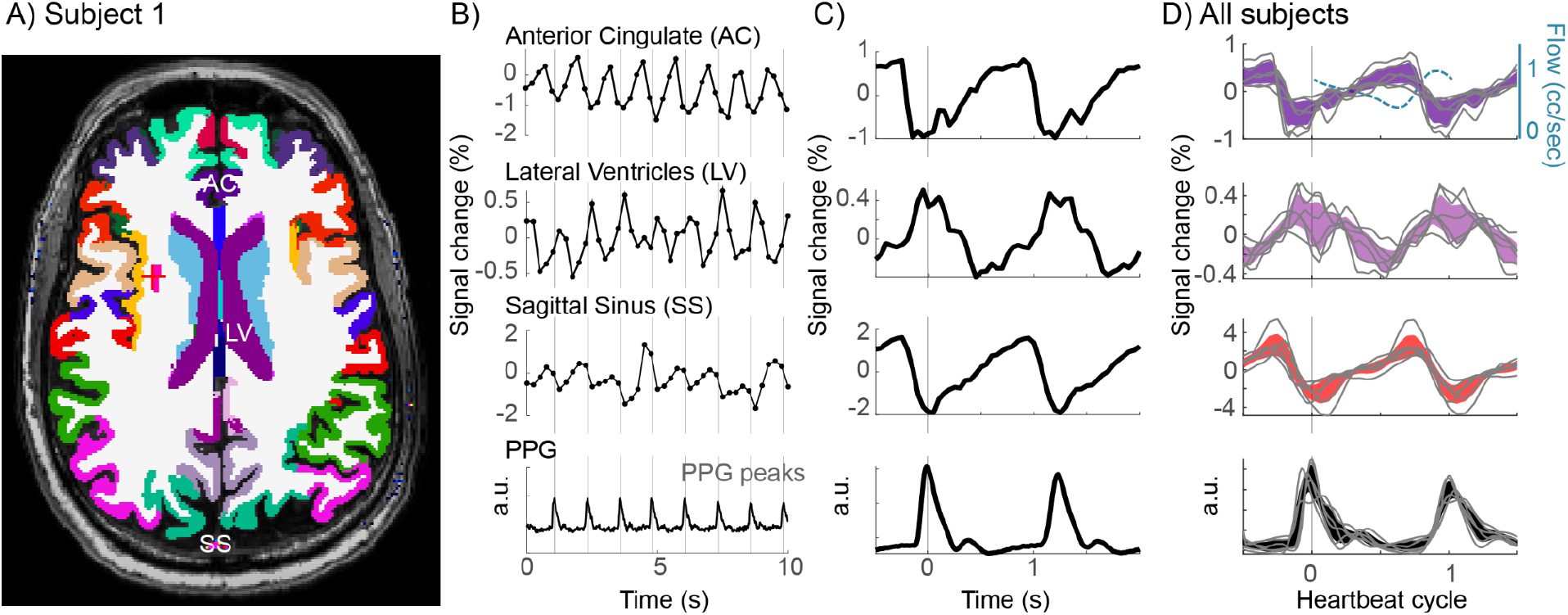
Reliable response shapes across subjects in several anatomical regions. **A)** Example slice with gray matter segmentation from Freesurfer, showing the anterior cingulate (AC), the lateral ventricle (LV) and the sagittal sinus (SS) segmented from the venogram. **B)** Functional MRI waveforms averaged across these three segmented areas, sampled every 250 ms. **C)** Aligning samples to each PPG peak allows a clear visualization of the waveform of the cardiac aligned responses within these regions of interest. **D)** Average curves for all 6 subjects when resampled as a function of the heartbeat cycle in each subject (mean +/ 2*SE). Gray lines show consistent shapes of cardiac aligned curves in each subject. The top panel also includes a curve of the typical flow speed (cc/sec) in the carotid artery (dashed light blue). The speed was measured using cine phase contrast MRI (Enzmann et al., 1994; Wagshul et al., 2011).

The cardiac aligned responses are not well described by a sinusoidal oscillation, but show a particular temporal asymmetry that matches known aspects of cerebral fluid circulation. The responses in the anterior cingulate and sagittal sinus drop sharply before the onset of the PPG peak, followed by a slow rise. We note that the PPG peak is measured on the thumb, and blood typically arrives in the brain before it arrives in the hand. Moverover, blood flow measurements in other studies from slices that include the carotid artery (Enzmann et al., 1994; Wagshul, Eide, & Madsen, 2011) show a sharp rise in the speed of flow related to systole that aligns with the sharp drop in our data (Figure 4D, top panel).

Both the anterior cingulate area, which is close to the anterior cerebral artery, and veins show a local minimum around the time of the PPG peak. The lateral ventricles show a signal change in the opposite direction, with a local maximum at the time of the PPG peak. The fact that an area close to a cerebral artery and the superior sagittal sinus behave in an opposite manner compared to CSF can be explained by the fact that the cranium is an enclosed space. Within this enclosed space, systole generates a rapid increase in blood pulsing into the cerebral arteries, which displaces CSF (Wagshul, Chen, Egnor, McCormack, & Roche, 2006; Wagshul et al., 2011). Whereas other methods have characterized these interactions across several seconds during sleep (Fultz et al., 2019), this method reveals these interactions at the timescale of a few hundred milliseconds within a heartbeat cycle.

### 3.3 Cerebral arteries influence the pulsatility in the gray matter

While Figure 4 illustrates reliable cardiac pulsations in the gray matter of the anterior cingulate, Figure 2 and Figure 3 show that many other gray matter regions do not have reliable cardiac pulsations. We therefore consider that the brain’s blood supply comes from three branches of the cerebral arteries; the posterior, anterior, and middle cerebral arteries (PCA, ACA and MCA). We hypothesize that the cardiac aligned responses in the gray matter will be large near the inputs to these three arteries. To measure this, we combined gray matter areas, segmented with Freesurfer, into three groups (Figure 5A,B) that are supplied by the three main cerebral arteries. Within each group, we assign an area to a subgroup depending on its distance from the main inputs. The rostral anterior cingulate and medial orbitofrontal cortex were closest to the main input from the ACA, the parahippocampal gyrus was closest to the PCA and the insular cortex and transverse temporal gyrus were closest to the MCA. Figure 5C shows that there is a decrease in pulsatility along the three main arteries: areas closest to the main arterial inputs have the largest cardiac averaged pulsations and more distant areas have smaller cardiac pulsations. This decrease in the amplitude of the cardiac aligned response with distance from the main input of each cerebral artery is reliably observed across subjects (Figure 5D). Moreover, the gray matter areas close to the main inputs of the three arteries also show similar response waveforms, with local minima before the PPG peak (time zero).

**Figure 5.**
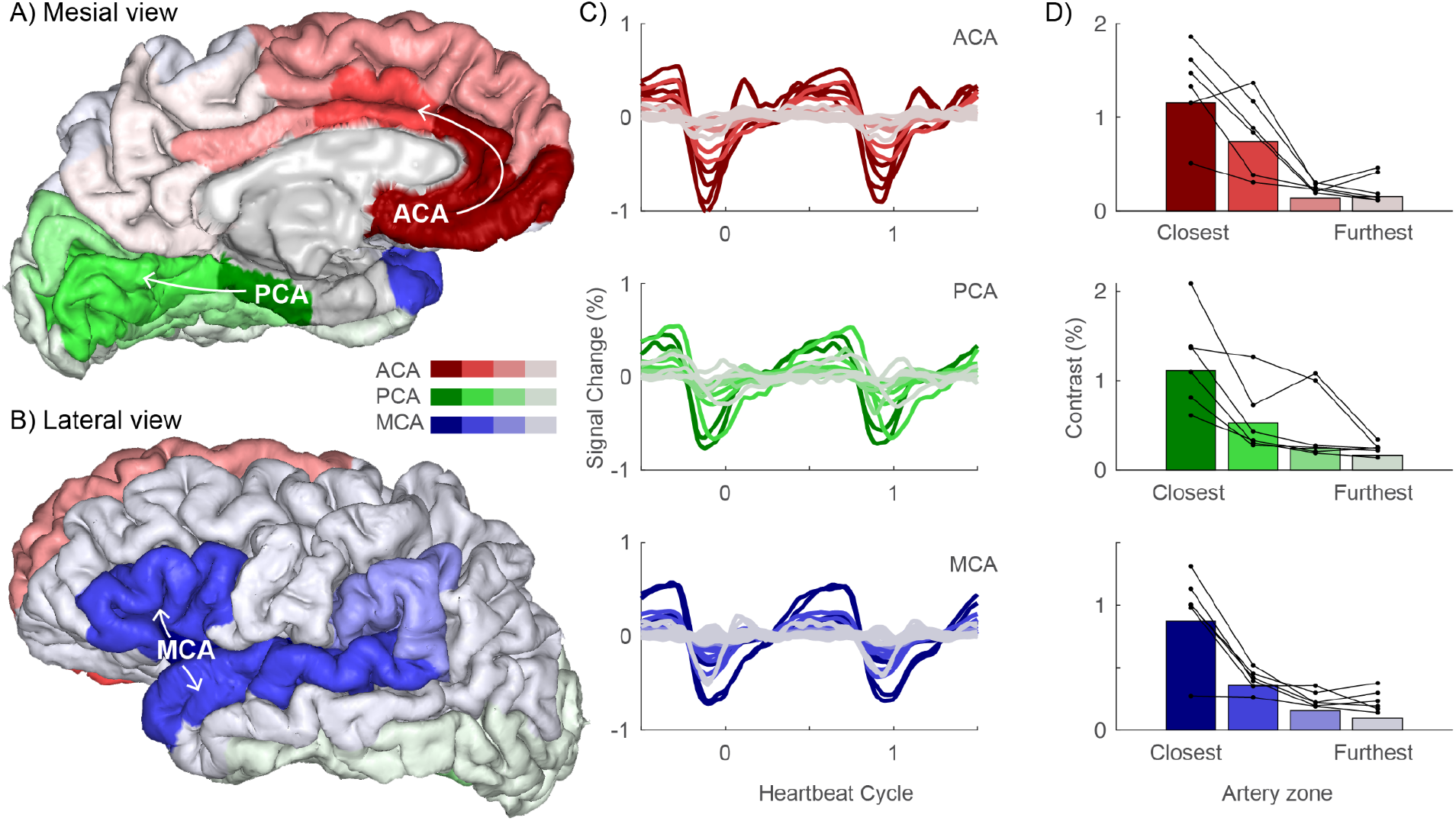
Response modulation in gray matter decreases with distance from the input to the main arteries. **A)** Mesial and **B)** lateral view of a cortex rendering segmented using Freesurfer and labeled according to the position on the three main arterial zones: the ACA (red), PCA (green) and MCA (blue). **C)** Average cardiac aligned responses across subjects during the heartbeat cycle in these three arterial zones. Lighter colors (gray) indicate areas further along each branch and typically show a decrease in modulation. **D)** The modulation amplitude within these areas averaged across subjects (bars), and for each subject (connected black lines/dots). There is one subject in each row of D) with the lowest contrast in the closest arterial zone; this is the same subject across all plots.

### 3.6 Timing of blood and CSF pulsations

The analysis of the fMRI time series shows that cranial spaces with blood and CSF have reliable cardiac aligned responses. Areas close to the inputs to the cerebral arteries show local minima close to the time of the PPG peak, and the ventricles show local maxima. The timing of the cardiac aligned responses provides information about the speed of the pulse pressure wave and can quantify delays between the arteries and veins. Pulse delays between arteries and veins could indicate the time that it takes for the cardiac pulse pressure waves to travel through the vascular system. Pulse wave velocity is related to arterial stiffening and pulse wave velocity measured between carotid and femoral arteries has been associated with an increased risk to develop dementia (Rouch et al., 2018). We therefore test whether the cardiac aligned responses allow us to map the timing of the pulse pressure wave.

To extract an image with the timing of the cardiac aligned responses, we perform a Singular Value Decomposition (SVD) to clean the waveforms (Figure 6A). We do this based on half the data, such that the validity of the model can be tested using the other half of the data. Two principal components explain over 68% of variance in the data (Figure 6B). The shape of these two components is highly consistent across subjects and allows extracting two group level canonical principal components (Figure 6C). The model with two canonical principal components fitted with cross validation (leave-one-subject-out) explains the data from the left out subject well (Figure 6D). The relative root mean squared error (*R*_*rmse*_) was smaller than 1 in 46.5%-83% of voxels (range across subjects), indicating that the model with two components explains the data from the left out subject better than within subject test-retest reliability. The leave-one-out cross validation suggests that the model will perform well in other subjects. A model based on these two canonical principal components describe various possible responses (Figure 6E and Supplemental Figure 2).

**Figure 6.**
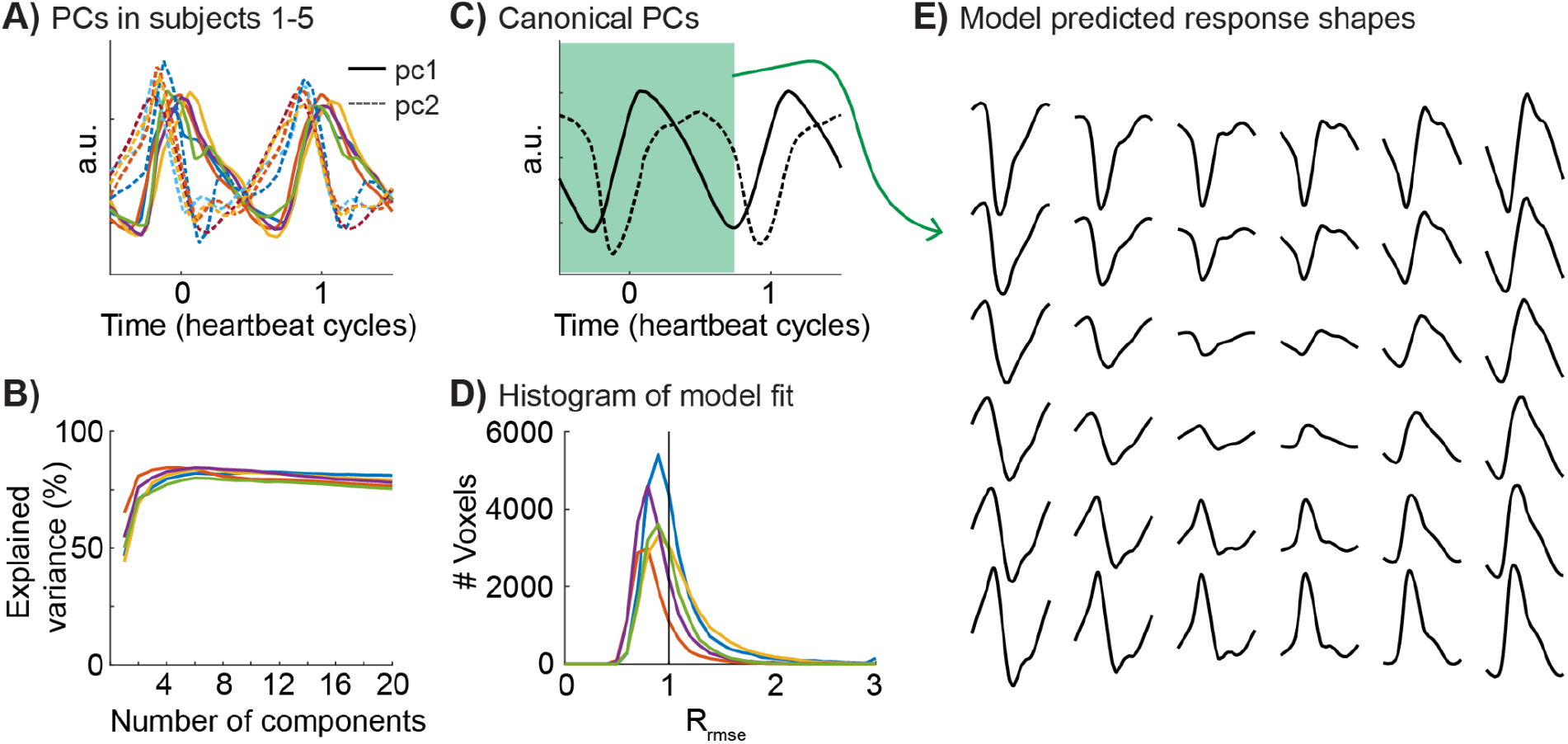
Model of the cardiac aligned responses. **A)** First, two principal components (PCs) for all five subjects plotted as a function of the heartbeat cycle. Time zero is the PPG peak. The first principal component is plotted in the solid line, the second component is plotted as a dashed line. **B)** The cross validated variance explained by an increasing number of principal components plotted for each subject. **C)** Canonical principal components across subjects as a function of heartbeat cycle. **D)** Histogram showing the distribution of the relative root mean square errors across voxels. A R_mse_<1 indicates that the model trained on 4 out of 5 subjects explained half of the data in the 5th subject better compared to the other half of the data from the same subject. Note that the expected value of a perfect model and Gaussian measurement noise lies around 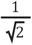. **E)** The model with these two principal components can predict various response shapes, the waveforms show time ranging from -0.50 to 0.66 of the heartbeat cycle (indicated in green in (C)).

This model based approach reduces noise and reliably characterizes the times of the local minima and maxima of the cardiac aligned responses. Figure 7A shows the average times of the local minima across subjects in terms of the percent of the cardiac cycle. Areas close to the main cerebral arteries such as the anterior cingulate cortex and the insula show pulses with a local minimum before the PPG peak (red/yellow). The superior sagittal sinus shows a later time on the local minimum right after the PPG peak (blue). Interestingly, these peaks are relatively close in time, separated only by about 20% of the cardiac cycle, which corresponds to ∼200 ms at a heart rate of 60 beats per minute.

**Figure 7.**
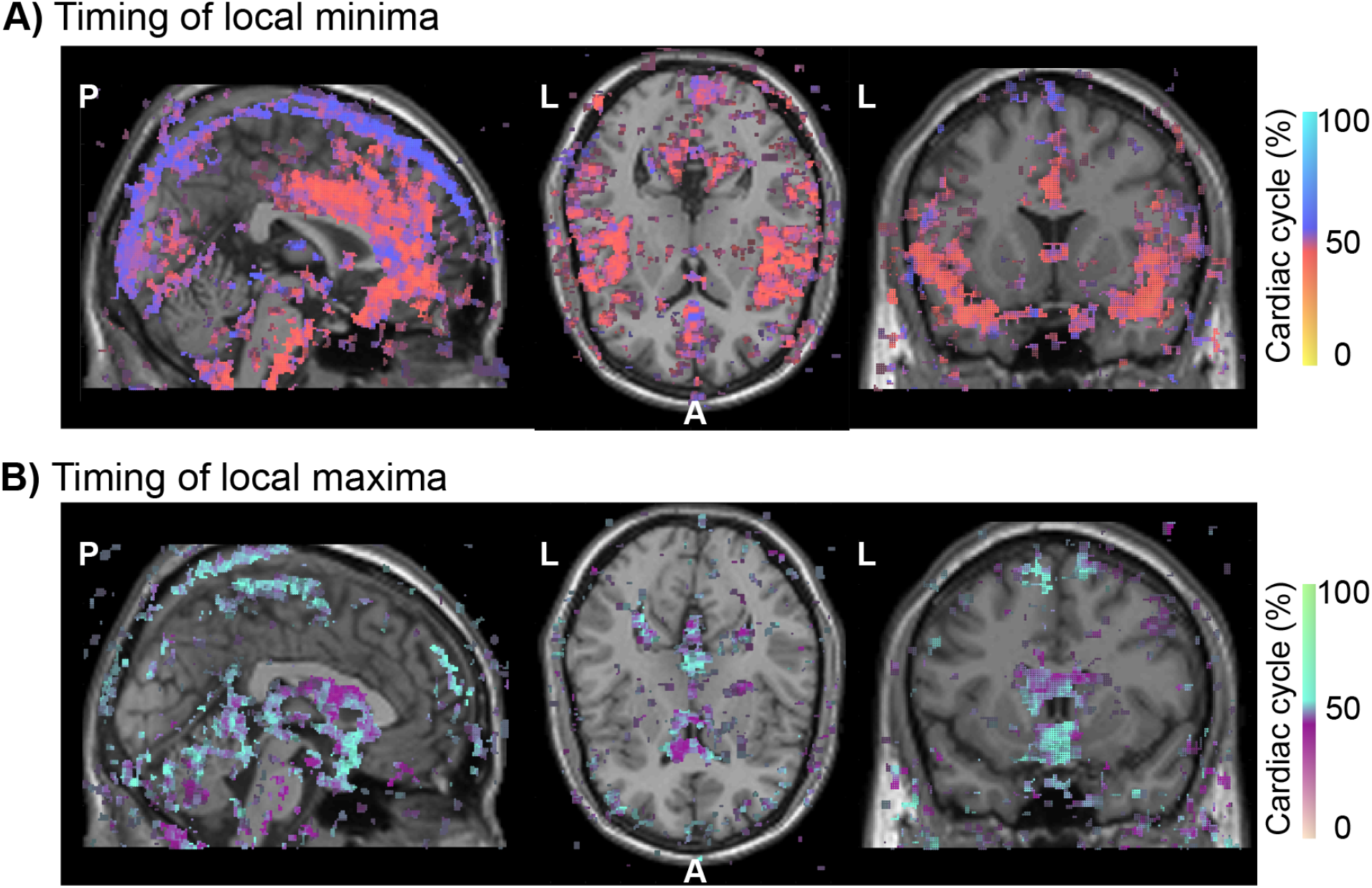
Spatial distribution of the timing of cardiac aligned responses in the cranium across the group. For each voxel in each subject, we calculated the peak time of the cardiac aligned response in terms of percentage of the cardiac cycle with the peak of the PPG at 50%. These timing maps were aligned in MNI152 space; in this way the average timing across the subjects could be calculated. **A)** The timing of the *local minima* for all voxels that had an average R^2^>0.3 across the subjects. Red colors indicate times before the PPG peak and blue colors indicate times after the PPG peak. **B)** The timing of the *local maxima* for all voxels that had an average R^2^>0.3 across the subjects. Purple colors indicate times before the PPG peak and cyan colors indicate times after the PPG peak.

Figure 7B shows the average times of the local maxima across subjects. Across subjects, the ventricles show the earliest times with a local maximum before the PPG peak (purple). Other areas with CSF, such as the subarachnoid cisterns and subarachnoid spaces under the superior sagittal sinus, show later pulses with a local maximum after the PPG peak (cyan). Similar as in the blood vessels, the difference in timing across the different CSF spaces is small, separated only by about 20% of the cardiac cycle.

This general pattern of response timing is also observed in individual subjects. Similar to the group average, individual subjects show early local minima in regions near the main three cerebral arteries and later local minima around the superior sagittal sinus (Figure 8A and 8B). Note that there are also significant local minima underneath the rendered cortex that likely correspond to the basilar and carotid arteries.

**Figure 8.**
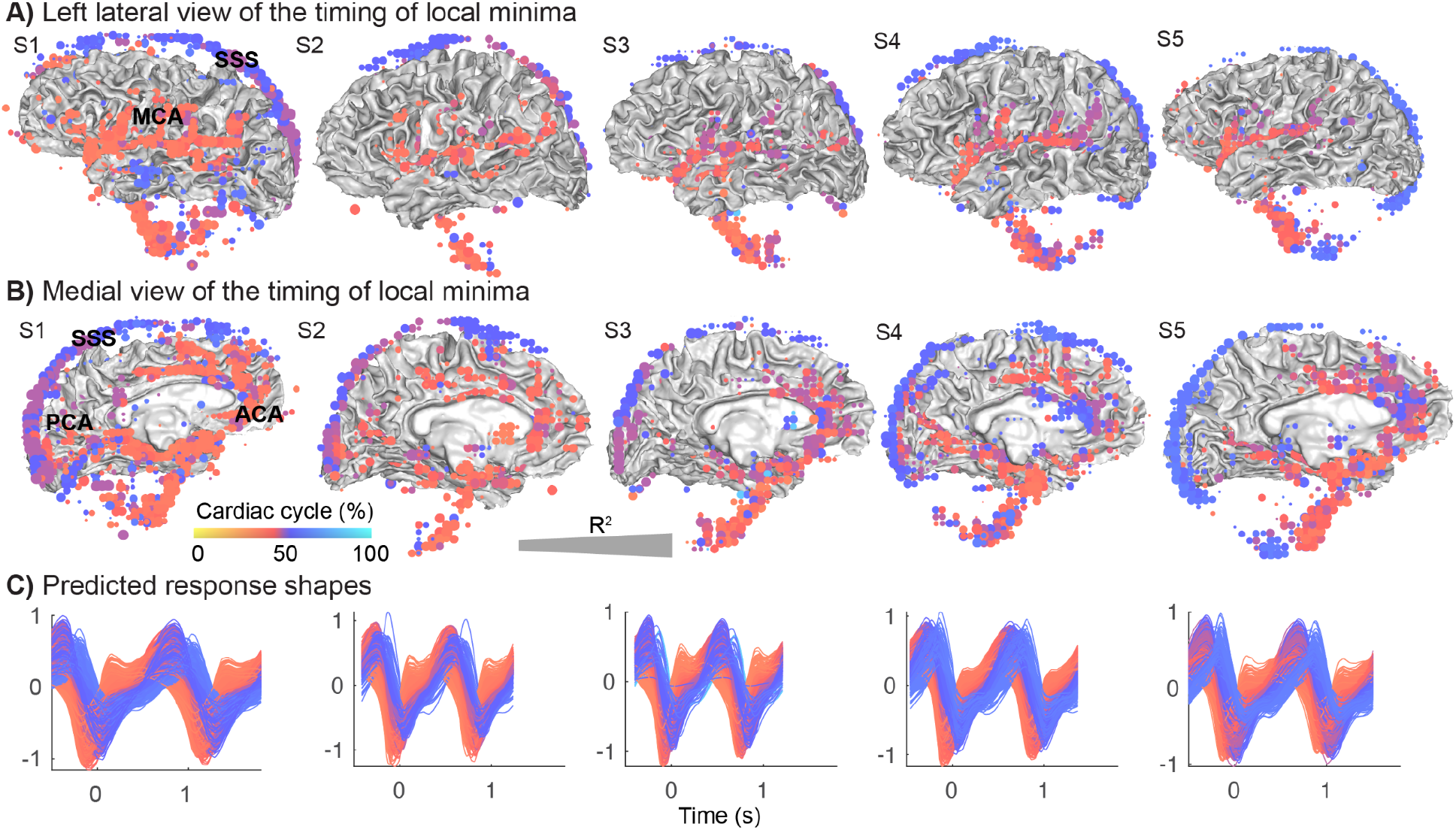
Distribution and timing of the local minima in the cranium of individual subjects. For all subjects (S1-S5) we calculated the timing of the local minima of the modeled response in terms of the percentage of the cardiac cycle for each voxel, with the peak in the PPG located at 50%. **A)** Left lateral view of times of local minima in voxels with reliable cardiac aligned responses (R^2^>0.5) were rendered on the gray/white matter surface. Red colors indicate an earlier minimum such as seen in areas close to the middle cerebral artery (MCA), and blue colors indicate a later minimum such as seen in the superior sagittal sinus (SSS). Larger dots indicate that voxels were more reliable (R^2^ scaled up to 1) **B)** Medial view of the times of the same local minima, red colors show earlier minima in areas close to the anterior cerebral artery (ACA), and posterior cerebral artery (PCA) and blue colors show later peaks such as those in the superior sagittal sinus (SSS). **C)** Predicted cardiac aligned response shapes for all displayed voxels as a function of time (s). Colors match the color of the voxels.

Similar to the average across the subjects, individual subjects also show early local maxima in areas near the ventricles and later local maxima in areas near the subarachnoid cisterns (Figure 9). Note that there are also significant local maxima underneath the rendered cortex that likely correspond to the CSF spaces near the fourth ventricle. However, there are inter-individual differences in the locations that show a local maximum in the reliable cardiac pulsation at later times. Note that the age range for the subjects was 24 to 63 years old.

**Figure 9.**
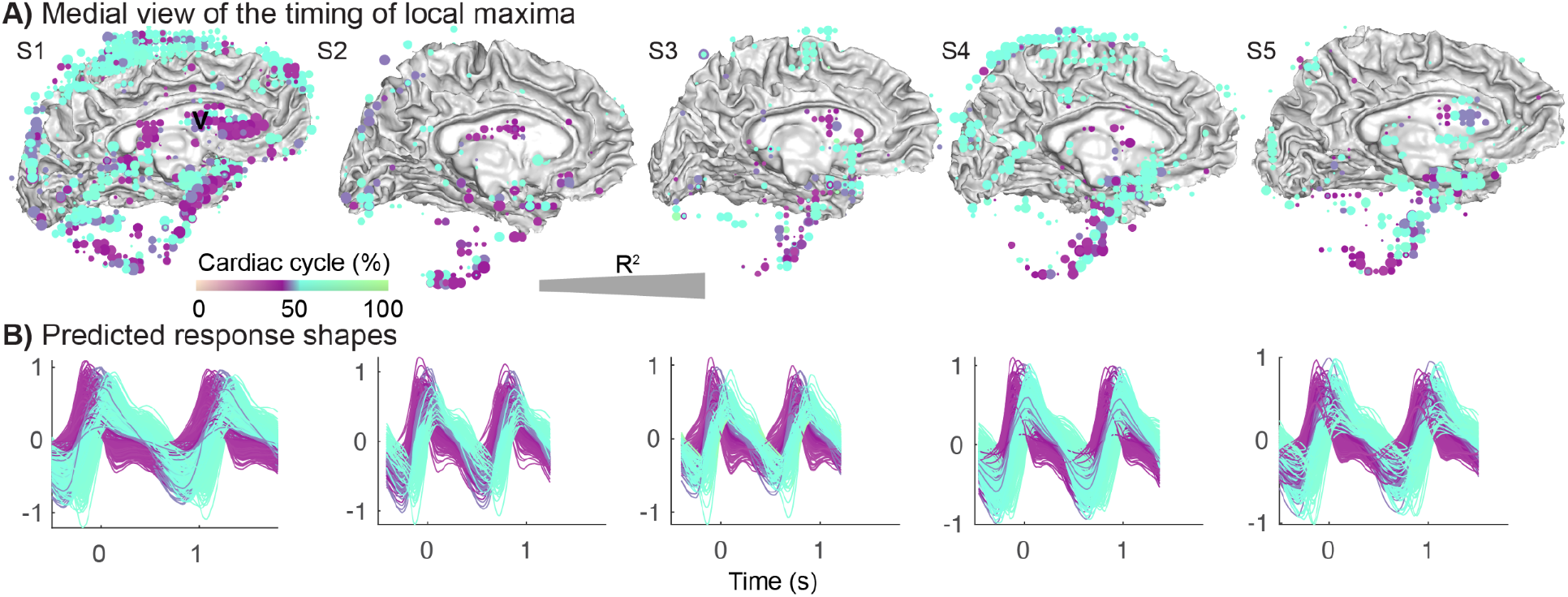
Distribution and timing of the local maxima in the cranium of individual subjects. For all subjects (S1-S5) we calculated the timing of the local maximum of the modeled response in terms of the percentage of the cardiac cycle for each voxel, with the peak in the PPG located at 50%. **A)** Medial view of times of local maxima in voxels with reliable cardiac aligned responses (R^2^>0.5) were rendered on the gray/white matter surface. Purple colors indicate peak times before the PPG peak and cyan colors indicate times after the PPG peak. **B)** Predicted cardiac aligned response shapes for all displayed voxels as a function of time (s). Colors match the color of the voxels.

### 3.7 Cardiac-gated resting state

In order to understand whether the characteristic blood and CSF pulsations could also be observed with a typical resting state fMRI scan, we analyze data from 5 subjects who were scanned in a different project (Hack et al., 2021). In these resting state scans, the echo time is slower and the repetition time is slower (SMS: TE = 11.6ms, FA = 48, TR = 250ms, resting state fMRI: TE = 27.5ms, FA = 77, TR = 2000ms), but each slice is still acquired within a short time of 44.4 ms. Each slice can be aligned to the PPG peak, and cardiac pulsations can be extracted. Similar to the fast SMS sequence, areas near main cerebral arteries, such as the left insula (Figure 10A) show large cardiac pulsations with a local minimum. Voxels in the lateral ventricles also show large cardiac pulsations, with a local maximum as in the SMS data (Figure 10B compared to Figure 4C). Similarly, the amplitude of the cardiac pulsations in the gray matter decreases with distance from the main 3 cerebral arteries (Figures 10C and 10D).

**Figure 10.**
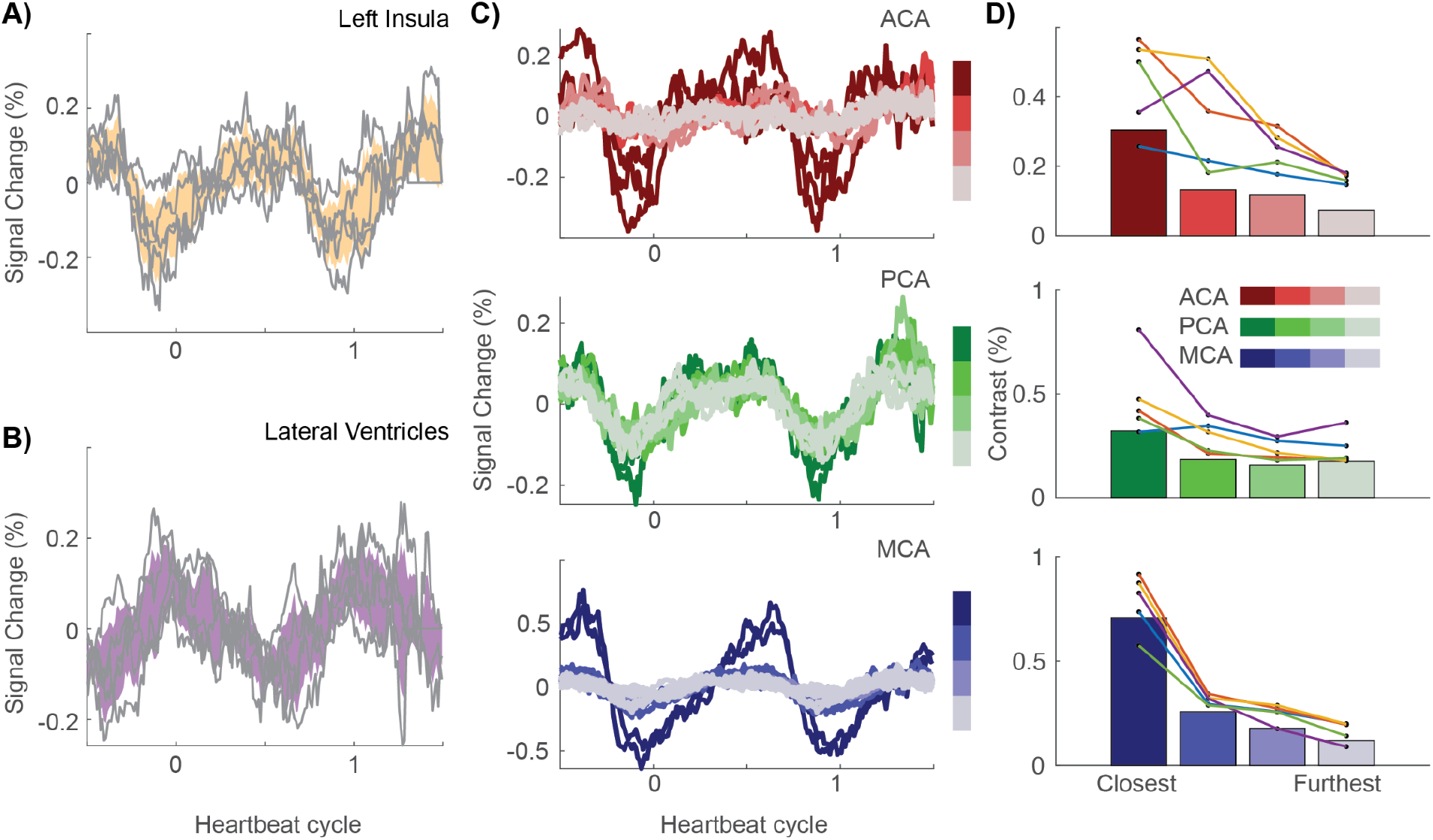
Cardiac aligned responses in resting state fMRI data. From a typical resting state fMRI scan, we extracted cardiac aligned responses. **A)** The mean cardiac aligned response in the insula for each subject in gray lines and average +/- 2 standard errors in yellow. **B)** The mean cardiac aligned response in the lateral ventricles for each subject in gray lines and average +/- 2 standard errors in purple. **C)** Average brain pulsations across subjects as a function of heartbeat cycle along these three arterial zoned ACA (red, top), PCA (green, middle) and MCA (blue, bottom). Lighter colors (gray) indicate areas further along each branch and typically show a decrease in pulsatility. **D)** Contrast within these areas averaged across subjects (bars), and for each subject (connected colored lines/dots).

The fact that cardiac alignment of the resting state fMRI results in similar response curves indicates that these responses are robust despite differences in scan sequences. Using the gradient echo signal equations (Hornak, 1996), we calculate that the SMS sequence has a higher sensitivity to changes in T1 than the resting state sequence, but slightly lower sensitivity to changes in T2*. Given the larger modulation measured using SMS, we believe the signal modulation is T1-weighted, likely carried by increases and decreases of the blood volume.

We note that the time in which one slice is acquired is comparable between the two methods: 50 ms for the SMS sequence and 44.4 ms for the resting state sequence. Much longer slice acquisition times would likely blur the cardiac pulsations. While cardiac aligned responses overall have similar characteristics, it has to be noted that there is more noise in the regular resting state acquisition compared to fast SMS acquisition. The resting state fMRI scan is about 2x longer and each slice is only acquired once every 2 seconds, while a slice is measured 4 times per second for the SMS data. Acquisition of fewer measurement time points per slice using an SMS sequence would also result in less reliable characterization of the cardiac pulsations (Supplemental Figure 3).

## 4. Discussion

The simultaneous multislice technique allows a rapid assessment of cardiac pulsations in arteries, veins and CSF. These pulsations produce a local minimum after systole in the fMRI signal in arteries and veins, and a local maximum in CSF spaces (Figures 2, 4, 5, 7, 8 and 9). The timing of these extrema matches known physiological dynamics in blood and CSF (Figure 4D). Early local minima are observed in gray matter regions near the three cerebral arteries (Figure 5) and later local minima are observed in the veins (Figure 7, 8). Early local maxima are observed in the ventricles and later local maxima are observed in the subarachnoid spaces (Figure 7, 9), with the latter having more spatial variability across subjects. An analysis of resting state fMRI data acquired with a conventional sequential slice acquisition shows that these physiological signatures are not unique to the SMS sequence (Figure 10), but are a general property of fMRI signals in blood and CSF.

### 4.1 Physiological basis of the cardiac-gated MR signal

The fMRI signal measures spin-coherence; a number of physiological properties jointly determine the signal level. By retrospective cardiac alignment, we obtain a temporal response that informs us about physiological signals that are principally driven by the beating heart. The signal analysis suggests that we are measuring two independent parameters: a signal amplitude and phase. Consequently, we have limited ability to make inferences about the many different factors that can give rise to the cardiac-gated fMRI signal (proton density, blood velocity, vessel stiffness) and how the signal depends on imaging parameters (TE, TR, FA, voxel size).

There has been some initial theory exploring the relationship between these factors for high field (7T) measurements that acquire a single fMRI slice (Bianciardi et al., 2016; Viessmann et al., 2017; Viessmann, Möller, & Jezzard, 2019). An important difference between these single slice 7T measurements and the whole brain measurements in our current study is that inflow and velocity influence the signals in a very different manner. When measuring a single slice, inflowing spins have not experienced prior excitation and signals follow a set of relatively well worked out equations (Bianciardi et al., 2016). The same equations may not apply to a whole brain acquisition. An example of the size of these effects can be found in a study measuring fMRI fluctuations on the timescale of seconds during sleep. The CSF in the fourth ventricle measured in the bottom slice of the acquisition showed large signal increases due to inflow at the same time as the fMRI signal in gray matter decreased (Fultz et al., 2019). These CSF signal increases were suppressed in slices that were not at the edge of the acquisition. While we characterize the temporal response shapes, it will be important to develop a better understanding of how whole brain fMRI signals are affected by different physiological parameters that fluctuate during the cardiac cycle such as blood or CSF volume, flow speed and partial voluming.

We find local minima in blood vessels around the time of the PPG peak. The local minimum is defined by a rapid decline when the cardiac pulse arrives, followed by a slow increase. The opposite pattern, with a local maximum, was observed in CSF spaces. These time courses were consistent between the fast SMS measurements and the resting state fMRI measurements. Another group, who defined a new whole brain Magnetic Resonance EncephaloGraphy (MREG) sequence showed similar dips in the MREG signal in the anterior cerebral artery following systole (Rajna et al., 2021). While a local minimum in blood spaces after systole is consistent between these studies, there is an important difference in our interpretations.

Rajna et al. assume that a local minimum in the MREG signal characterizes the pulse pressure wave in all fluid spaces, both blood and CSF. In contrast, we find that the pulse pressure wave induces a local minimum in blood but a local maximum in CSF spaces. Our assumption is in line with cranial fluid dynamics that show the cardiac pulse has opposite effects on blood and CSF (Wagshul et al., 2006). During the heartbeat, blood moves in and occupies a larger volume, and CSF moves out and occupies a smaller volume (Beggs, 2014). This explains why the time course in blood and CSF would be inverted.

### 4.2 Complementary nature of various MRI based measurements of cardiac pulsations

The pulsatile flow during the cardiac cycle is accompanied by several mechanical and physiological factors that can be assessed in various complementary ways. Cine MRI measurements encoded for displacements have shown that some brain areas move about 0.1 mm during the cardiac cycle (Soellinger, Rutz, Kozerke, & Boesiger, 2009). Amplified MRI analyses represent these cardiac-gated signals as movements of the ventricles, blood vessels and at gray or white matter boundaries (Abderezaei et al., 2020; Kolipaka, White, & Ehman, 2021; Terem et al., 2018).

Further understanding the relation between the pulsatile motion and fluid dynamics is important. Pial blood vessels are surrounded by CSF (Iliff et al., 2013). One study measured cardiac cycle induced arterial wall motions (Mestre et al., 2018) using particle tracking velocimetry and two photon imaging through a sealed cranial window in mice. Arterial diameters were about 10μm and increased with about 0.1μm during the cardiac cycle. Average arterial diameters did not change during hypertension, but the typical vessel wall speeds were increased in hypertension, in particular in more distal arteries. These hypertension related increases in wall motion were further accompanied by a reduced forward CSF flow in surrounding CSF spaces. The use of complementary techniques may further elucidate different physiological factors noninvasively in humans in neurological and neuropsychiatric diseases.

### 4.3 Retrospective cardiac alignment reveals temporal features of the cardiac response

While our data have a relatively low spatial resolution with 4mm isotropic voxels, our measurements have the advantage that with a brief fMRI scan, they reveal detailed temporal profiles of the pulsations in blood and CSF spaces. We retrospectively aligned the measured signals to the PPG peak, which allowed us to observe that the cardiac pulsations are not sinusoidal, with a steep slope when the pulse arrives followed by a shallow slope on return. This asymmetry in the pulsatile waveform has been related to fluid dynamics. For example, the slope and amplitude of the pulse pressure wave may be related to intracranial compliance, with a less compliant tissue resulting in a steeper slope (Wagshul et al., 2011). A study in an older population used a high resolution 4D flow MRI measurement with velocity encoding sensitive to arterial blood flow speed to study the relation between the slope of the cardiac pulse in the cerebral arteries and episodic memory (Vikner et al., 2021). They found that steeper systolic onsets correlated with poorer episodic memory performance. The ability of our technique to extract the shape of the cardiac pulse, including slope and width, in both blood vessels and CSF spaces may help further elucidate these types of effects.

### 4.4 Implications for resting state fMRI analyses

Previous studies have emphasized the importance of understanding the effects of cardiac pulsations on resting state fMRI networks (Bayrak et al., 2021; Chen et al., 2020; Shmueli et al., 2007). One study examined the effects of changes in heart rate on the fMRI response (Chang et al., 2009) and identified a heart rate response function that evolves over a few seconds with changes in heart rate. Although we are interested in these heartbeat related signals, these resting state studies emphasize the removal of cardiac effects (Glover, Li, & Ress, 2000; Hu, Le, Parrish, & Erhard, 1995). These effects can be significant and hard to remove because cardiac pulsations influence fMRI signals in brain regions within a second, but fMRI data are typically sampled every one to two seconds. This undersampling results in aliasing of heartbeat related signals. Two studies have shown that faster measurements can reduce the size of these unwanted signals (Huotari et al., 2019; Jahanian et al., 2019).

There are many large resting state fMRI datasets with TRs of 1-2 seconds. Our slow fMRI analyses demonstrate that such measurements can be retrospectively aligned when a pulse oximetry measurement is available. The cardiac aligned responses could be used in a forward manner to correct for unwanted heartbeat driven fluctuations in the resting state signal.

### 4.5 Conclusion

Cardiovascular mechanisms are essential for healthy cognitive and affective function. Neurological diseases and aging affect the way in which the cardiac pulse affects the brain’s fluid dynamics. To understand whether fMRI data can help assess the spatial distribution and temporal delays of the cardiac pulsations, and conversely, how cardiac pulsations affect the fMRI signal, we collected data with a fast SMS sequence and retrospectively aligned the measurements to heartbeats. Cardiac aligned responses reveal the combined impact of the cardiac pulse pressure wave and blood flow on the fMRI signal during the cardiac cycle. Identifying the typical range of responses in the healthy may help us identify atypical responses in neurological and psychiatric diseases.

## 5. Acknowledgements

The authors thank Aviv Mezer, Ariel Rokem, Rosemary Le and Dominic Konrad for their help in collecting the data. This work was supported by a Stanford CNI innovation grant.

## 6. Glossary of terms

ACA: Anterior Cerebral Artery
CSF: CerebroSpinal Fluid
FA: Flip Angle
fMRI: functional Magnetic Resonance Imaging
MCA: Middle Cerebral Artery
MNI: MOntreal Neurological Institute
MREG: Magnetic Resonance Encephalography
PC: Principal Component
PCA: Posterior Cerebral Artery
PPG: PhotoPlethysmoGraphy
SMS: Simultaneous MultiSlice
SVD: Singular Value Decomposition
TE: Echo Time
TR: Repetition Time

## Supplemental Materials

**Supplemental Figure 1.**
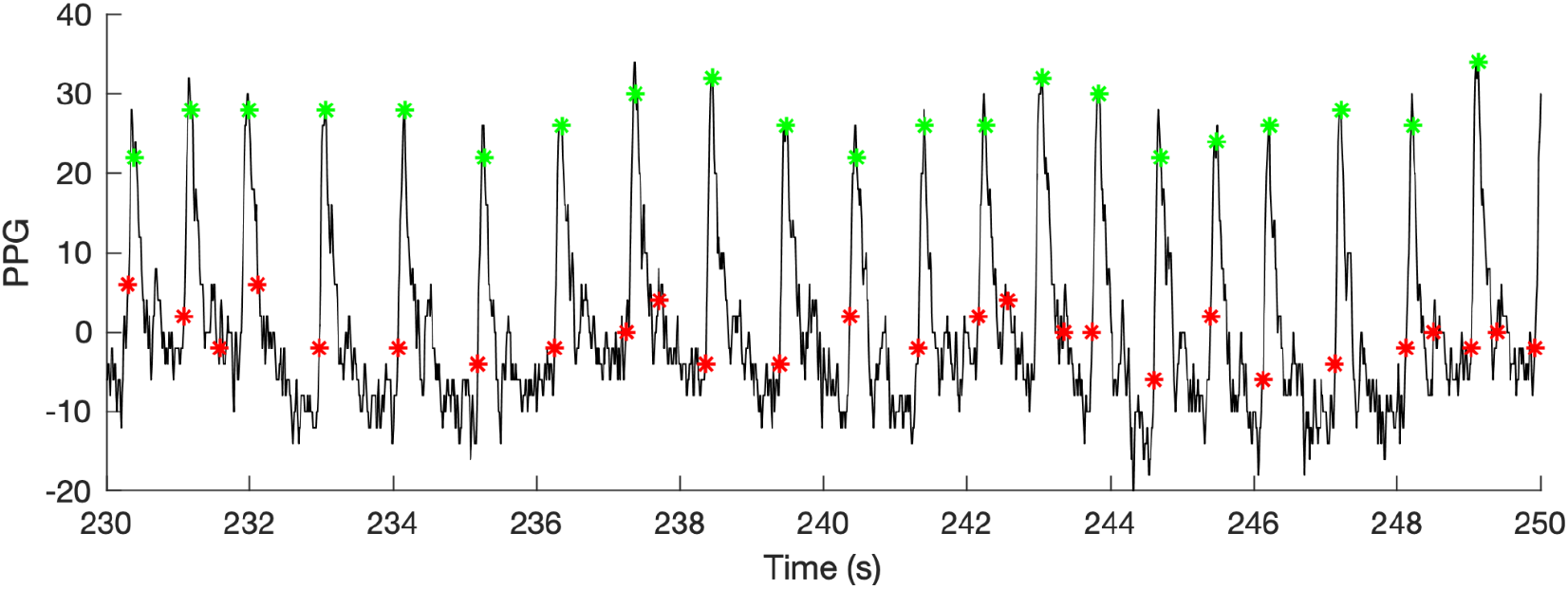
Detection of heartbeats in the raw PPG signal shown as a function of time (seconds). Red dots show the heart beats detected by the GE scanner software in real time, note that there are several false positive peaks detected and a few peaks are missed. Green dots show the heart beats detected by our algorithm (section 2.3). Peaks are detected in the low pass filtered signal, and green dots sometimes appear a few samples to the left or right of the peak, but there were fewer false positives.

**Supplemental Figure 2.**
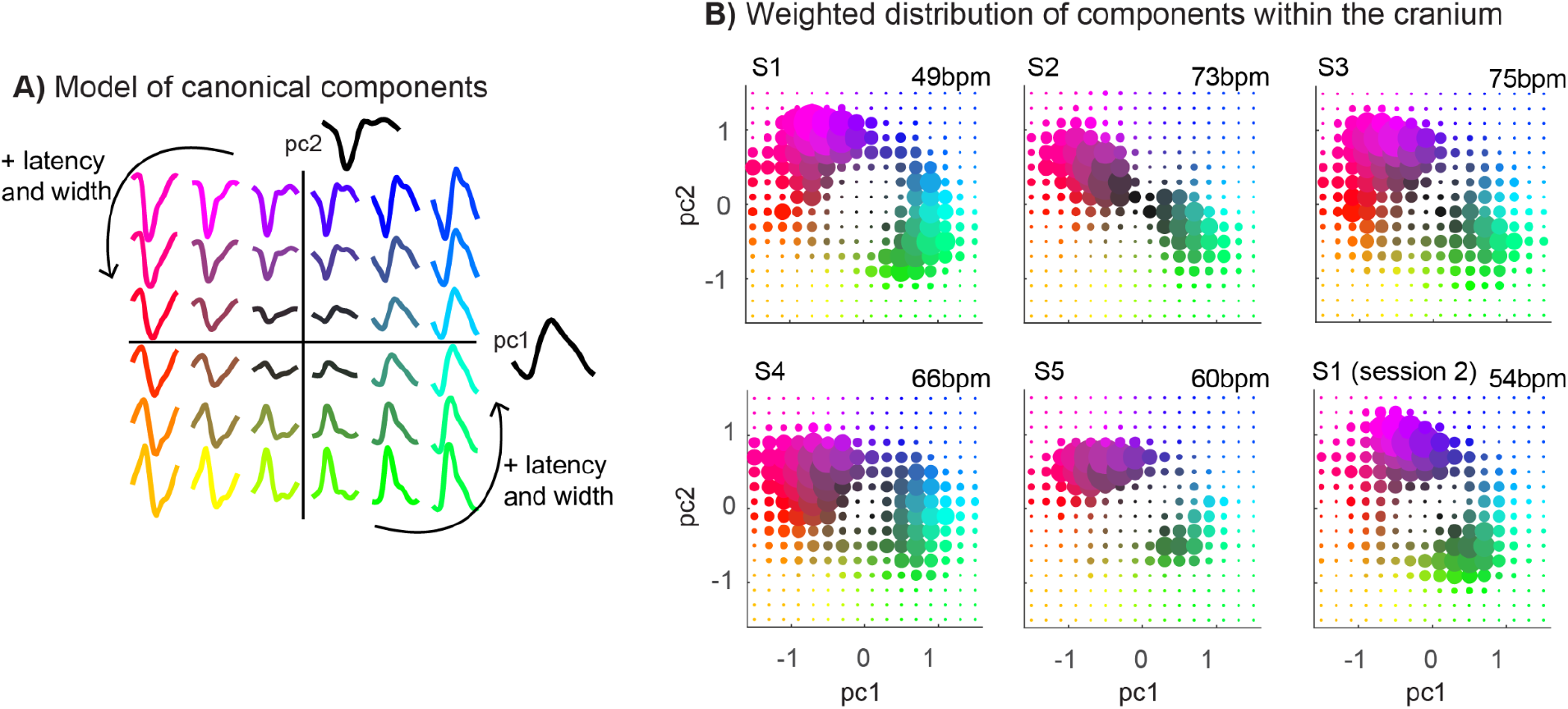
Modeling the cardiac aligned responses. **A)** Canonical components (pc1 and pc2) across the subjects can be combined with various weights to predict different cardiac aligned response shapes. **D)** Fitting these two canonical components in each subject shows that component weights cluster into two groups with low pc1, high pc2 weights (purple - red) and with high pc1, low pc2 weights (green - blue). The dot size indicates the number of voxels with component weights. Only highly reliable voxels with R^2^>0.7 were included.

**Supplementary Figure 3.**
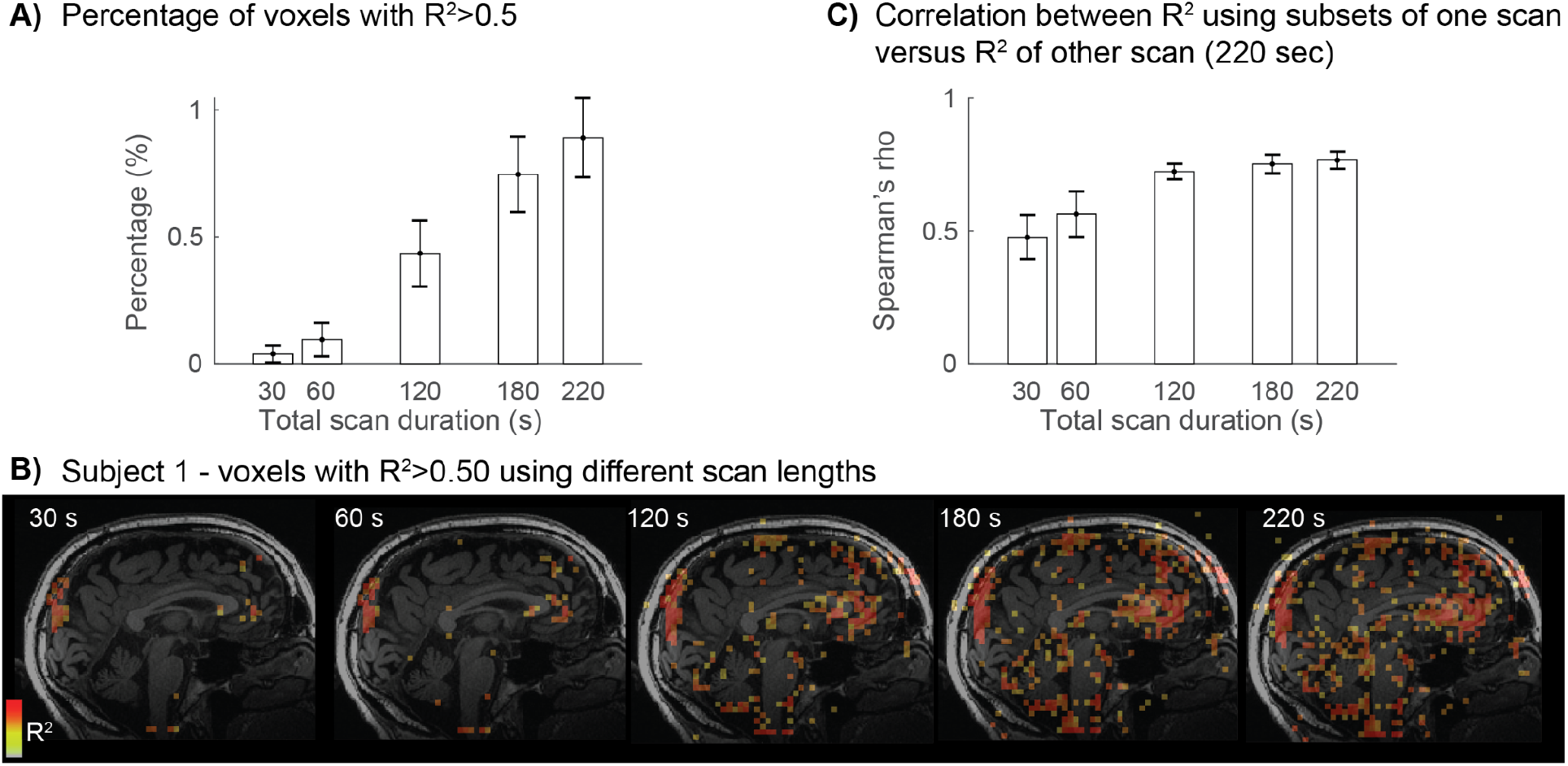
Investigation of different scan durations. **A)** As an indication of the number of areas where we calculated the cardiac pulsations with less data. The total scan duration was 220 seconds, and we also used 30, 60, 120, 180 sec of data to calculate the percentage of voxels with reliable heart beat aligned responses. Reliable responses were considered those voxels with an R^2^>0.5 between even and odd responses. Bars show the average across subjects +/- one standard deviation across the subjects. **B)** Example of the voxels with reliable cardiac pulsations (R^2^>0.5) calculated based on different scan length overlaid on a T1 image in one subject. It can be seen that with scan durations shorter than 120 seconds, many regions are missed at this threshold. **C)** To understand whether a shorter scan duration result in different patterns of reliable cardiac pulsations, or whether the reliability simply decreases while the overall pattern remains the same, we calculated the spatial correlation (Spearman’s rho) between the R^2^ calculated based on a subsample of one scan versus the R^2^ calculated on a separate 220 sec long scan. The y-axis shows the average across subjects +/- one standard deviation. This suggests that with shorter scan durations, similar spatial patterns of reliable cardiac pulsations will be found when scanning for 120 seconds or longer, but cardiac aligned response shapes will be more reliable with longer scan durations (calculated up to 220 sec). We note that longer scan durations may increase chances of subject motion and lower reliability.

## Notes

### Competing Interest Statement

The authors have declared no competing interest.

